# Co-occurring alterations in the RAS-MAPK pathway limit response to MET inhibitor treatment in *MET* exon 14 skipping mutation positive lung cancer

**DOI:** 10.1101/374181

**Authors:** Julia K. Rotow, Philippe Gui, Wei Wu, Victoria M. Raymond, Richard B. Lanman, Frederic J. Kaye, Nir Peled, Ferran Fece de la Cruz, Brandon Nadres, Ryan B. Corcoran, Iwei Yeh, Boris C. Bastian, Petr Starostik, Kimberly Newsom, Victor R Olivas, Alexander M. Wolff, James S. Fraser, Eric A. Collisson, Caroline E. McCoach, D. Ross Camidge, Jose Pacheco, Lyudmila Bazhenova, Tianhong Li, Trever G. Bivona, Collin M. Blakely

**Affiliations:** Department of Medicine, University of California, San Francisco; Helen Diller Family Comprehensive Cancer Center, University of California, San Francisco; Guardant Health, Inc; Department of Medicine, University of Florida; Soroka Medical Center, Ben-Gurion University, Beer-Sheva, Israel; Massachusetts General Hospital Cancer Center and Department of Medicine, Harvard Medical School; Departments of Dermatology and Pathology, and Clinical Cancer Genomics Laboratory, University of California, San Francisco; Department of Pathology Immunology and Laboratory Medicine, University of Florida; Department of Bioengineering and Therapeutic Sciences, University of California, San Francisco; University of Colorado; University of California, San Diego; Departiment of Internal Medicine, University of California, Davis

## Abstract

**PURPOSE:** While patients with advanced-stage non-small cell lung cancers (NSCLCs) harboring *MET* exon 14 skipping mutations (*METex*14) often benefit from MET tyrosine kinase inhibitor (TKI) treatment, clinical benefit is limited by primary and acquired drug resistance. The molecular basis for this resistance remains incompletely understood.

**METHODS:** Targeted sequencing analysis was performed on cell-free circulating tumor DNA obtained from 289 patients with advanced-stage *METex*14-mutated NSCLC.

**RESULTS:** Prominent co-occurring RAS-MAPK pathway gene alterations (e.g. in *KRAS, NF1*) were detected in NSCLCs with *METex*14 skipping alterations as compared to *EGFR*-mutated NSCLCs. There was an association between decreased MET TKI treatment response and RAS-MAPK pathway co-occurring alterations. In a preclinical model expressing a canonical *METex*14 mutation, KRAS overexpression or NF1 downregulation hyperactivated MAPK signaling to promote MET TKI resistance. This resistance was overcome by co-treatment with crizotinib and the MEK inhibitor trametinib.

**CONCLUSION:** Our study provides a genomic landscape of co-occurring alterations in advanced-stage *METex*14-mutated NSCLC and suggests a potential combination therapy strategy targeting MAPK pathway signaling to enhance clinical outcomes.

## Introduction

Somatic *MET* mutations leading to splicing-mediated loss of exon 14 and subsequent MET overexpression are an emerging therapeutic target present in 2-4% of lung adenocarcinomas (1, 2). MET tyrosine kinase inhibitor (TKI) treatment was associated with improved overall survival in a retrospective study of patients with *METex*14-mutated NSCLC (3). In ongoing prospective studies, response rates of 32%, to the multi-kinase inhibitor crizotinib, 41%, to the MET TKI tepotinib, and up to 72% for treatment-naïve patients to the MET TKI capmatinib have been reported (4–6). Crizotinib has recently received FDA breakthrough designation for use in the treatment of *METex*14-mutated NSCLC.

While MET remains an attractive therapeutic target, both primary and acquired resistance limit the long-term survival of patients with *METex*14-mutated NSCLC. Second-site *MET* mutations and downstream signaling reactivation via acquired *KRAS* amplification have been reported at acquired resistance to MET TKI therapy, and may inform treatment decisions (7–10). By contrast, the mechanisms mediating both primary MET TKI resistance and tumor cell persistence during initial MET TKI treatment remain largely undefined, as does the full landscape of alterations promoting acquired resistance.

Recent studies show that advanced-stage NSCLCs often harbor multiple oncogenic alterations, which may impact response to targeted therapies (11–13). Analysis of cell-free circulating tumor DNA (cfDNA) utilizing next-generation sequencing (NGS) provides one avenue to describe the genomic landscape within a cancer patient and offers the potential to capture genomic changes reflecting heterogeneity across distinct metastatic tumor sites (11, 14). A more detailed understanding of the mutational landscape that co-exists with oncogenic *MET* alterations in NSCLC may facilitate an improved understanding of the determinants of MET TKI response and identify rational polytherapy strategies to improve clinical outcomes.

Here, we describe the spectrum of co-occurring genomic alterations observed within the cfDNA of patients with *METex*14-mutated, advanced-stage NSCLC and identify prominent co-alteration of RAS pathway genes as a contributing factor to disease progression.

## Results

### Co-occurring genomic alterations are common in advanced-stage METex14-mutated lung cancer

We analyzed a cohort of 289 patients with advanced-stage NSCLC who had a *METex*14 mutation identified upon cfDNA analysis of 68 cancer-relevant genes using a clinically validated assay (Guardant360). This is the largest reported cohort to-date describing the genomic landscape of advanced-stage **METex*14-mutated* NSCLC. We evaluated the frequency with which *METex*14 mutations co-occur with other cancer-associated mutations (Supplemental Table 1) (11, 15). Synonymous mutations and those with predicted neutral or unknown functional impact were excluded, as previously described (11), as were mutations previously associated with clonal hematopoiesis (16). 86.5% of samples contained co-occurring genomic alterations, with a mean of 2.74 alterations per sample (range 0-22), in addition to the *METex*14 mutation. The most commonly altered genes, co-occurring with the *METex*14 mutation in at least 10% of patients, were *TP53* (49.5% of patients), *EGFR* (16.3%), *NF1 (neurofibromatosis-1)* (15.6%), *BRAF* (10.7%), and *CDK4* (10.4%). Additional *MET* gene alterations were also common in this patient population, in which 9.3% of patients had co-occurring *MET* copy number gain and 12.5% had a co-occurring second-site *MET* mutation (Figure 1A).

**Figure 1.**
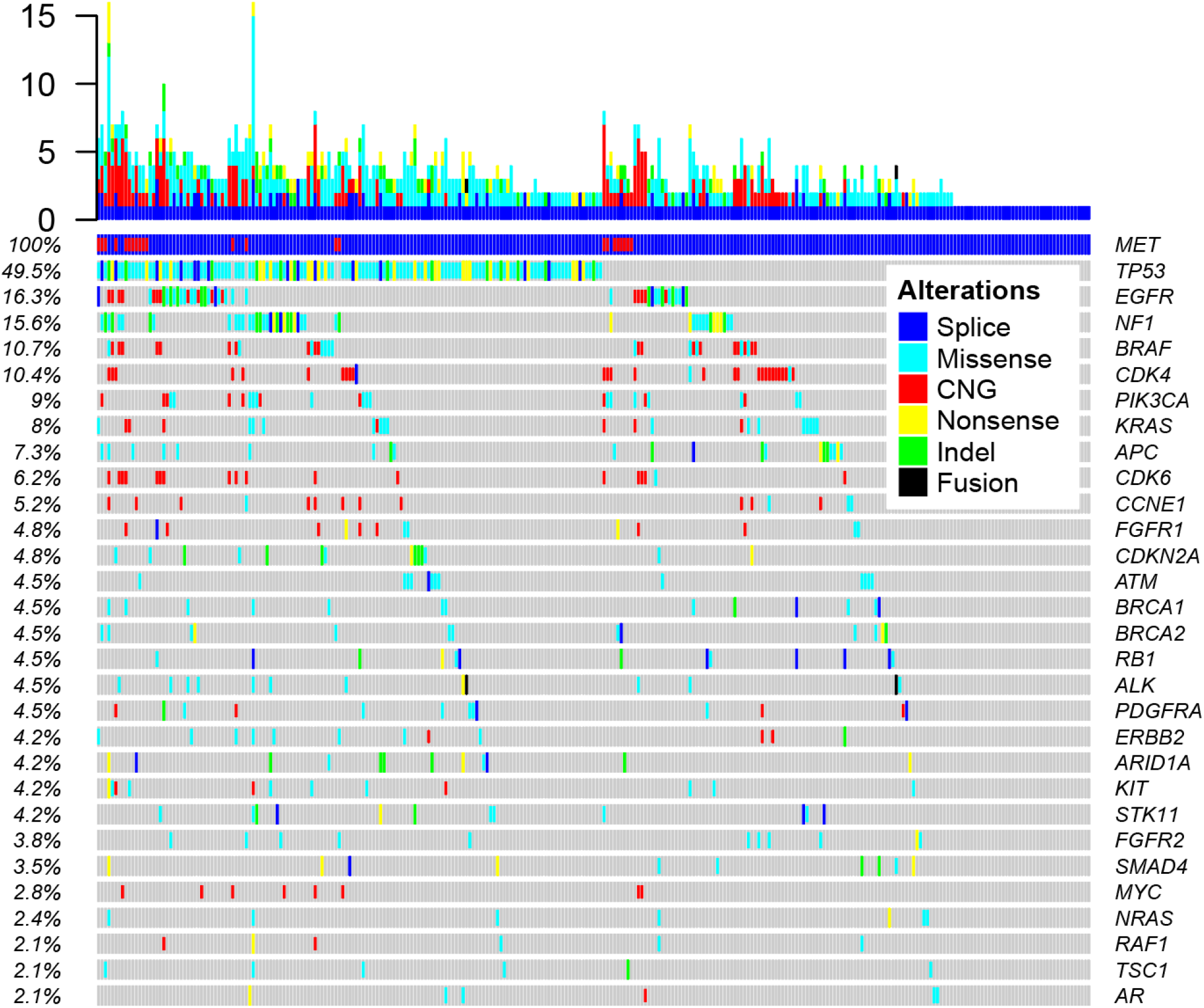
Co-occurring genomic alterations are common in non-small cell lung cancer (NSCLC) with a *MET* exon 14 mutation. **A.** Distribution of co-occurring genomic alterations in a targeted list of cancer-associated genes (Supplemental Table 2) as detected by cfDNA in 332 samples from 289 patients with advanced-stage *METex*14-mutated NSCLC. Results are filtered to exclude synonymous variants, variants predicted to result in an unknown or neutral function impact (via COSMIC v83, GENIE, ClinVar, and mutation assessor prediction algorithms) and mutations previously reported as associated with clonal hematopoiesis. Only genomic alterations occurring at a 2% or greater frequency are displayed. CNG, copy number gain.

Seventeen of 34 second-site *MET* mutations were located in the tyrosine kinase domain. All identified *MET* second-site mutations, regardless of predicted functional impact, were included in this analysis. Of these, fourteen (G1163R, L1195V/F, F1200I, D1228H/N/Y, and Y1230H/S) were located at residues previously associated with MET TKI resistance (7–9, 17–19). Nine of these occurred in patients with known prior MET TKI exposure and five in patients with unknown prior treatment history. The remaining three mutations (H1094Y, R1336W, and I1084L) occurred in patients without available prior treatment history. While the *MET* H1094Y mutation is known to lead to MET activation (20), the effects of the other two *MET* mutations remain uncharacterized.

The co-occurrence of other established oncogenic driver alterations (*KRAS, EGFR, ALK, ROS-1, BRAF*) was uncommon except for the presence of activating *KRAS* mutations in 5.2% of patients (G12C/D/S/V 3.5%, G13C 1%, Q22K 0.3%, and Q61H 0.3%), canonical EGFR-activating mutations in 3.5% (del19 3.1%, L858R 0.7%, T790M 2%), and an *ALK* gene rearrangement in 0.7%. In addition, a *HER2* exon 20 insertion was detected in one patient. In a patient with known clinical outcomes data, with both *EGFR* del19/EGFR T790M mutations and a *METex*14 mutation, there was partial response (RECIST 1.1) to treatment with the EGFR TKI osimertinib, which lasted 13.8 months. While in the overall dataset the *METex*14 mutation detected was predominantly clonal (80.6% of samples), in samples with a detectable co-occurring oncogenic driver alteration the *METex*14 alteration was more likely to be subclonal (21.4% of samples clonal, p-value <0.0001). The converse was also true; co-occurring established oncogenic driver mutations that were detected were more likely to be clonal than the other detected co-occurring genomic alterations (68.6% versus 46.3%, p-value = 0.0144) (Supplemental Figure S1).

### RAS pathway alterations are more common in METex14-mutated NSCLC than in EGFR-mutated NSCLC

To understand how the genomic landscape in advanced-stage *METex*14-mutated NSCLC compares to NSCLCs with a different targetable oncogenic driver mutation associated with high upfront response rates to TKI therapy, we used identical cfDNA profiling to compare a cohort of patients with known *METex*14-mutated NSCLC to a previously unpublished, independent cohort of 1653 samples from 1489 patients with advanced-stage EGFR-mutant (del19, L858R) NSCLC (Supplemental Table 1). This comparison demonstrated differential frequency of co-occurring genomic alterations in 17 genes (Figure 2A). In the *METex*14-mutated cohort, co-occurring alterations in *NF1, CDK4, STK11, ALK, KRAS, ATM, CDKN2A, NRAS, TSC1*, and *ESR1* were more commonly identified. In the EGFR-mutated cohort, co-occurring alterations in *AR, ERBB2, CCNE1, PIK3CA, BRAF, CTNNB1*, and *MYC* were more commonly identified, independently validating our prior findings identifying these as common co-occurring alterations in EGFR-mutated NSCLC (11). Among the 10 genes with a greater frequency of co-occurring genomic alteration in patients with a *METex*14 mutation, three (*NF1, KRAS*, and *NRAS*) are key components of the RAS-MAPK signaling pathway.

**Figure 2.**
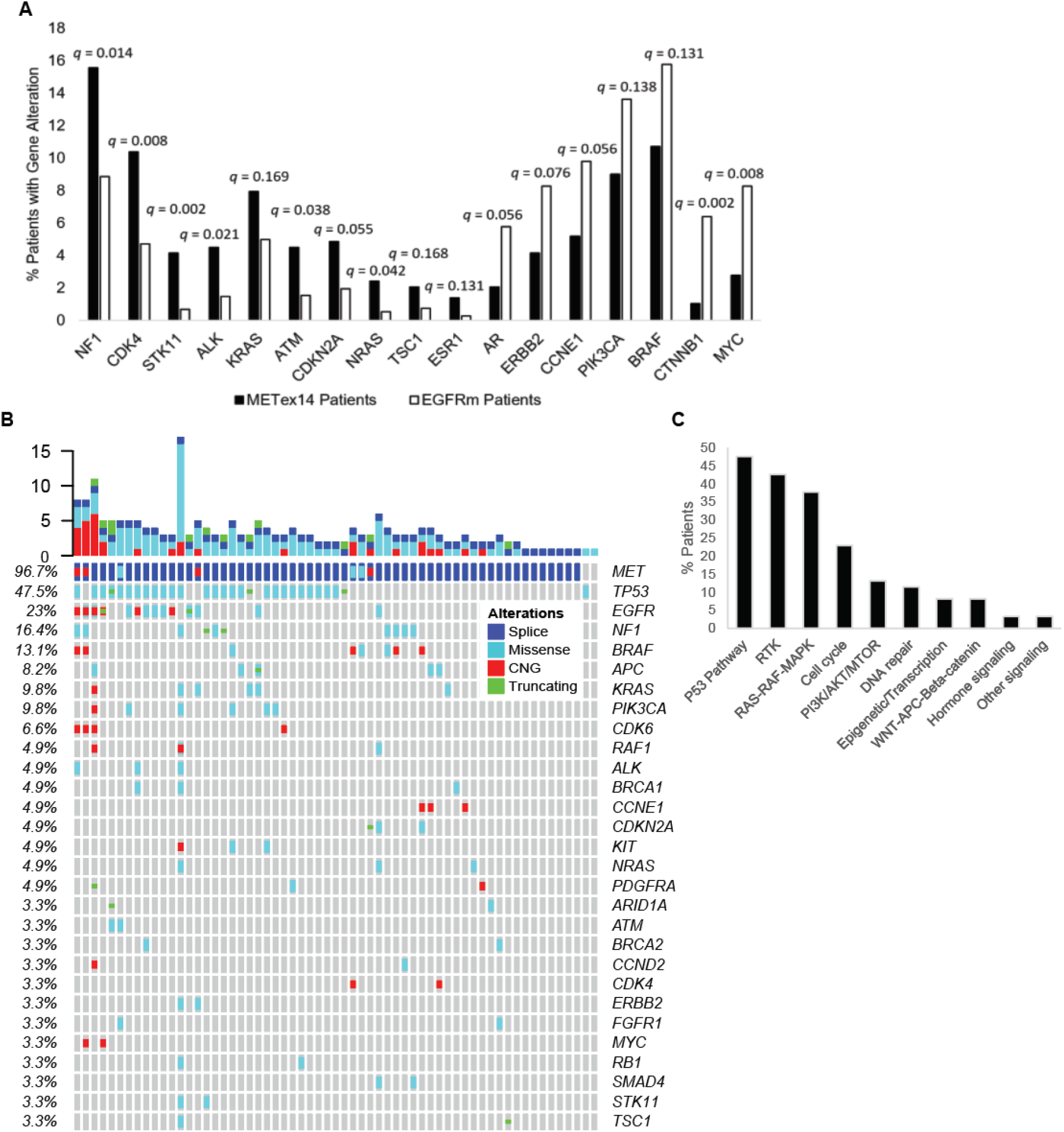
RAS-MAPK pathway alterations are common in *METex*14-mutated NSCLC. **A.** Comparative frequency of genomic alterations as measured by cfDNA in a cohort of advanced-stage *METex*14 NSCLC patients (n=289) compared to an independent cohort of patients (n=1489) with a known canonical *EGFR* activating mutation (*EGFR* exon 19 deletions, *EGFR* L858R). Gene alterations with a statistically significant difference are displayed. q-values calculated by two-tailed Fisher’s exact test with Benjamini-Hochberg multiple hypothesis testing for false discovery rate < 0.2. **B.** Spectrum of gene alterations detectable by cfDNA in the subset of *METex*14-mutated NSCLC patients (n=61) without prior MET TKI treatment. Genomic alterations with a frequency of greater than 2% displayed. **C.** Genomic alterations detectable by cfDNA in the same patient cohort classified by functional category. CNG, copy number gain.

A propensity towards genomic alterations promoting downstream hyperactivation of the RAS pathway may favor primary resistance to TKI therapy and help explain the comparatively lower TKI response rates in *METex*14-mutated NSCLC. Examination of the subset of cfDNA samples obtained from patients with *METex*14-mutated NSCLC prior to known MET TKI treatment (n=61) demonstrated high rates of genomic alterations capable of promoting RAS-MAPK pathway activation (Figures 2B, 2C). When compared to a previously published cohort (11) of cfDNA samples from TKI-naïve patients with EGFR-mutated NSCLC (n=20), RAS-MAPK pathway alterations that were detected before treatment remained more common in *METex*14-mutated NSCLC (35.2% vs 10%, p-value = 0.042).

### Both MET second-site mutations and RAS pathway alterations are newly detectable following MET TKI treatment

We identified twelve patients with cfDNA obtained following treatment with crizotinib and an available matched sample obtained prior to known crizotinib exposure (Figure 3A, B, Supplemental Table 3). Details regarding one patient (patient #5) in this dataset have previously been published by other groups (7, 9).

**Figure 3.**
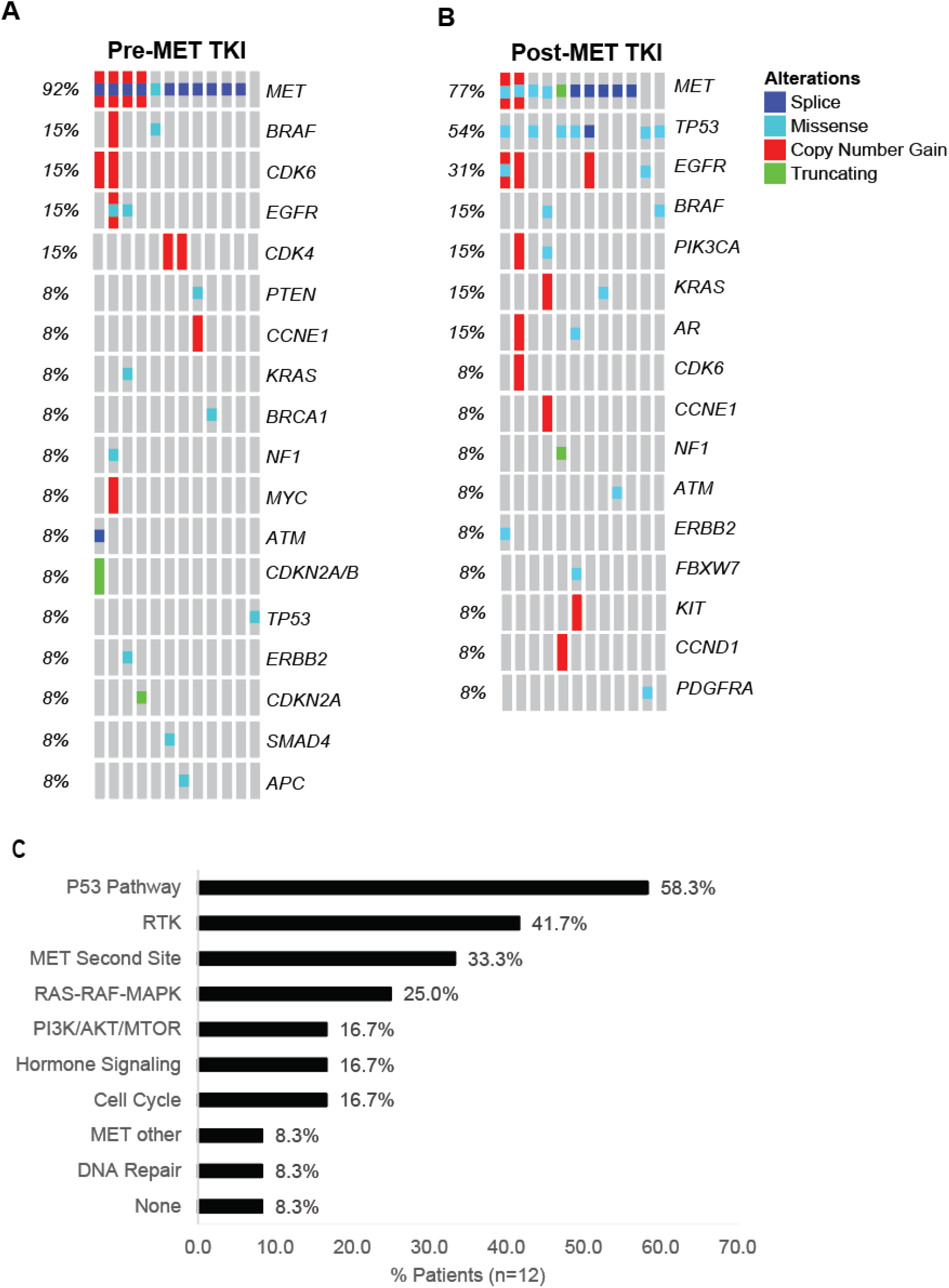
*MET* second-site and bypass pathway alterations are newly detectable after MET TKI treatment. Oncoprints show detectable genomic alterations by in paired samples from 12 patients, obtained either **(A)** before known MET TKI exposure or **(B)** after known MET TKI exposure. All samples reflect the results of cfDNA analysis (Guardant360), with the exception of three patients for whom only tissue NGS was available pre-MET TKI. Genes not common to the cfDNA panel were excluded from analysis. Full details available in Supplemental Table 3. **C.** Spectrum of newly detectable genomic alterations following MET TKI exposure by functional category.

Newly detectable *MET* second-site mutations (Y1230H/S, D1228H/N, F1200I, L1195V) were detected in four of twelve patients following MET TKI treatment (Figure 3C, Supplemental Table 3). While some of these mutations have been reported at acquired MET TKI resistance (7–9, 17–19), the *MET* F1200I has not yet been described in a patient sample. Identification of specific acquired second-site mutations at resistance to MET TKI therapy may inform treatment decisions. MET TKIs can be classified as type I TKIs (e.g. crizotinib, capmatinib) or type II TKIs (e.g. cabozantinib) based on the kinase domain conformation to which they bind and differ in their activity against second-site MET mutations. For example, the *MET* Y1230X and D1228X mutations develop at resistance to type I MET TKIs and may predict response to subsequent type II MET TKI treatment (7–9, 17–19). Conversely, the *MET* L1195V mutation has been reported at resistance to type II MET TKI treatment (7).

The development of acquired *MET* F1200 mutations as a resistance mechanism to MET TKIs has previously been predicted in a preclinical drug resistance screen, in which *MET* F1200 mutations were the dominant resistance mechanism to a type II MET TKI and were observed, though less common, at resistance to a type I MET TKI (21). Molecular modeling studies suggest that *MET* F1200I alters the conformation of the kinase domain such that it interferes with both the binding of type II MET TKIs within the DFG-out binding pocket, and to a lesser extent, may promote type I MET TKI resistance through disruption of an autoinhibitory MET conformation (Supplemental Figure S2) (7). Moreover, the F1200 residue is conserved across multiple tyrosine kinases, including ALK, ROS1, NTRK, and ABL, in which mutations at the corresponding residue have been linked to TKI resistance (Supplemental Table 4) (21–27).

In contrast, parallel and downstream pathway alterations with the potential to provide alternative input for RAS-MAPK pathway signaling were newly detectable in eight of twelve patients following MET TKI treatment as compared to cfDNA samples obtained prior to MET TKI treatment (Figure 3C, Supplemental Table 3). In addition to two patients with *KRAS* amplification (Figure 4), which was recently implicated in MET TKI resistance (10), an *NF1* frameshift mutation, and copy number gain in *EGFR* and *KIT* were identified. In one patient with acquired KRAS amplification, addition of the MEK inhibitor trametinib to crizotinib treatment rapidly decreased detectable circulating tumor cfDNA. In the second patient with acquired *KRAS* amplification, an activating *KRAS* G12D mutation was also present within a progressing lesion on crizotinib. Further details regarding the two patients with acquired *KRAS* amplification are presented in Supplemental Figure 3.

**Figure 4.**
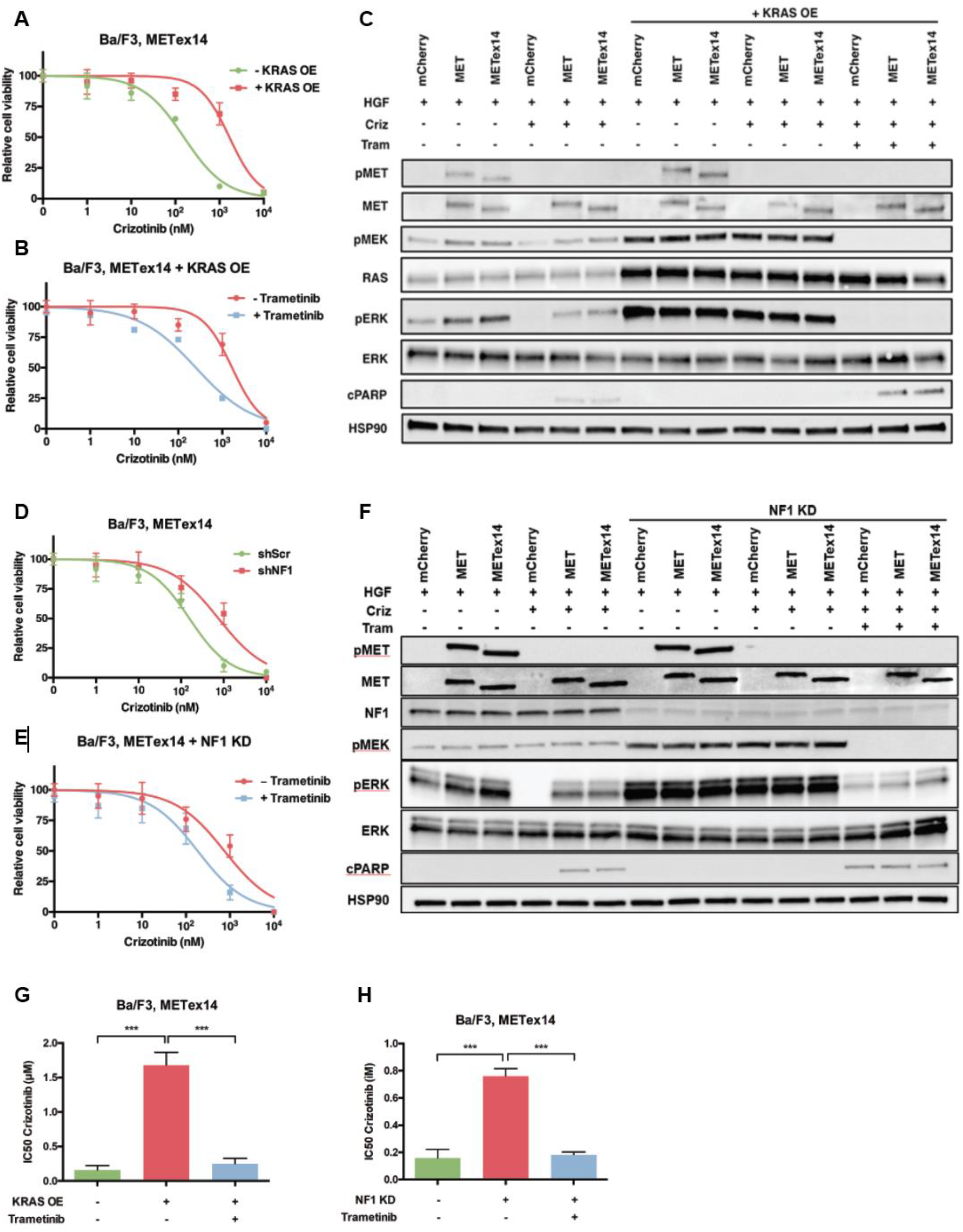
*KRAS* overexpression or *NF1* downregulation promotes MET TKI resistance that is overcome by combined crizotinib and trametinib polytherapy in *METex*14-mutated preclinical models. **A.** Cell viability curves demonstrating a shift in the half maximal inhibitory concentration (IC50) to crizotinib with overexpression of wild type *KRAS* in *MET* exon 14-mutant expressing Ba/F3 cells. The cells were grown in culture with HGF supplementation (50 ng/mL). **B.** Cell viability curve for crizotinib treatment in Ba/F3 cells with both *METex*14 expression and wild-type *KRAS* overexpression (KRAS OE), in the setting of treatment with trametinib at 0.01 μM and supplementation with HGF 50 ng/mL. The KRAS OE, trametinib negative curves in panels A and B reflect the same experimental data, displayed on two graphs for clarity. **C.** Cell viability curves for crizotinib-treated MET exon 14-mutant expressing Ba/F3 cells with either NF1 knockdown (shNF1) or a negative control scrambled shRNA (shScr). **D.** Cell viability curves for crizotinib-treated MET exon-14 mutant expressing Ba/F3 cells with NF1 knockdown, with and without addition of 0.01 μM trametinib. The shNF1, trametinib negative curves in panels C and D reflect the same data displayed on two graphs for clarity. **E.** Ba/F3 cells with stable expression of wild type MET, MET with an exon 14 skipping mutation (*METex*14), or mCherry control were treated with 50 ng/mL HGF with or without 24 hours treatment with crizotinib at 0.1 μM and/or trametinib (0.01 μM). Treatment with crizotinib inhibits MET phosphorylation and inhibits downstream Erk phosphorylation, with associated increase in apoptosis as measured by cleaved PARP (cPARP). KRAS overexpression (KRAS OE) restored downstream Erk phosphorylation and reduced cleaved PARP, despite crizotinib treatment. Addition of trametinib inhibited Erk phosphorylation and increased cleaved PARP, consistent with induction of apoptosis, despite the presence of KRAS overexpression. **F.** NF1 knockdown (NF1 KD) restores downstream Erk phosphorylation and decreases cleaved PARP in crizotinib-treated tumor cells. The addition of trametinib reduced Erk phosphorylation and restored PARP cleavage consistent with induction of apoptosis despite the presence of NF1. **G.** Summary graph of IC50 to crizotinib with trametinib with and without KRAS overexpression. **H.** IC50s of crizotinib with trametinib with or without NF1 downregulation. *** p-value < 0.001 by student’s t-test.

### MEK inhibition overcomes crizotinib resistance induced by RAS-MAPK pathway alterations

The clinical data suggested that preexisting or acquired genomic changes leading to RAS-MAPK pathway activation (e.g. in *KRAS, NF1*) may limit response and induce resistance to MET TKI treatment in *METex*14-mutated NSCLC. In one patient with *KRAS* amplification detectable at acquired resistance to crizotinib treatment, combination treatment off-label with both crizotinib (250 mg po BID) and trametinib (2 mg po daily) resulted in rapid loss of both detectable *METex*14 and *KRAS* amplification by cfDNA suggestive of treatment response. However, despite molecular evidence of tumor response this combination therapy was poorly tolerated in the context of overall clinical decline, with fatigue, fluid retention, and diarrhea. Despite dose reduction to trametinib 2 mg every other day, the patient expired before radiographic response assessment (Figure S3).

To assess the functional impact of RAS-MAPK pathway hyperactivation on sensitivity to MET TKI treatment, we engineered a new Ba/F3 cell-based system. The IL-3 dependent Ba/F3 cell line, while not of epithelial origin, is an established system to assess oncogenic capacity and putative drug resistance mechanisms (28, 29). Stable expression of human *METex*14 in Ba/F3 cells in the presence of the MET ligand human HGF (Hepatocyte Growth Factor) induced IL-3-independent growth and increased downstream Erk phosphorylation compared to expression of wild type *MET* (Figure 4D, 4F). *METex*14-mutant expressing cells were sensitive to treatment with crizotinib, as measured by reduced cell growth in standard cell viability assays (Figure S4).

Overexpression of wild type *KRAS* or knockdown of wild type *NF1* in *METex*14-expressing Ba/F3 cells induced resistance to crizotinib (Figure 4A, 4D). Overexpression of wild type *KRAS* increased the IC50 (inhibitory concentration of drug that decreases cell viability to 50%) to crizotinib from 0.16 μM to 1.68 μM (p-value < 0.001) (Figure 4A, 4G) and *NF1* downregulation increased the IC50 to crizotinib from 0.16 μM to 0.75 μM (p-value < 0.001) (Figure 4D, 4H). Treatment of cells harboring both *METex*14 and *KRAS* overexpression with the combination of crizotinib and the MEK inhibitor trametinib in order to block both MET signaling and downstream MAPK pathway signaling restored sensitivity to treatment (crizotinib IC50 of 1.68 μM in the absence of trametinib versus 0.25 μM with trametinib co-treatment, *p*-value < 0.001) (Figure 4B, 4G). Similarly, co-treatment of cells harboring both *METex14* and *NF1* downregulation with both trametinib and crizotinib restored sensitivity to treatment (crizotinib monotherapy IC50 0.75 μM versus 0.18 μM upon addition of trametinib, *p*-value < 0.001 (Figure 4E, 4H). The selected trametinib dose modestly reduced but did not eliminate cell growth in the absence of crizotinib (Figure S4). Immunoblotting demonstrated sustained Erk phosphorylation despite crizotinib treatment in samples with *KRAS* overexpression or *NF1* downregulation, which was abrogated by the addition of trametinib (Figure 4C, 4F). In both genomic contexts, combination treatment was associated with increased levels of cleaved PARP, indicative of apoptosis, which was absent with monotherapy (Figure 4C, 4F).

## Discussion

The challenge of therapeutic resistance in patients harboring *METex*14 mutations is of increasing clinical relevance given the emergence of MET-targeted therapies into the clinic. While off-label use of MET TKI therapy has demonstrated clinical activity, objective reported response rates of approximately 30-70% as reported in early studies (4–6) are generally lower than those seen in response to TKI treatment in NSCLC driven by other canonical oncogenes (EGFR, ALK), where response rates greater than 80% have been reported (30, 31).

RAS-MAPK pathway hyperactivation has an established role in promoting resistance to EGFR, ALK, BRAF, and ROS1 targeted therapies via diverse mechanisms including *KRAS* amplification and *KRAS, BRAF*, and *NF1* mutations (32–40). While *KRAS* amplification has also recently been reported at acquired MET TKI resistance in *METex*14-mutated NSCLC (10), the broader role of compensatory genomic events providing signaling pathway re-activation remains less well-understood in *METex*14-mutated NSCLC. Here, we describe frequent co-occurring RAS-MAPK pathway alterations in *METex*14-mutated NSCLC as compared to EGFR-mutated NSCLC. RAS-MAPK pathway alterations were detected even in TKI-naïve patients; our preclinical and clinical data suggest these co-occurring alterations may promote resistance to MET TKI therapy. More specifically, RAS-MAPK pathway alterations when present as co-occurring genomic events in *METex*14-mutated NSCLC prior to MET TKI treatment may induce not only primary resistance but also contribute to tumor cell persistence, thus limiting response magnitude and potentially duration during initial treatment. This notion is supported by our findings in the Ba/F3 preclinical system we engineered. We also describe the spectrum and relative frequencies of newly detectable genomic alterations following MET TKI treatment as measured by cfDNA, which included alterations within the RAS-MAPK pathway or within RTKs upstream of the RAS-MAPK pathway (41) in two thirds of patients. While our reported dataset is limited by lack of complete clinical outcomes data, it highlights the importance of developing future patient cohorts incorporating outcomes data to link the understanding of the genomic landscape to treatment response and prognosis, particularly for those patients with less common or emerging driver mutations.

Clinically, while many *MET* second-site mutations acquired during type I MET TKI (e.g. crizotinib) treatment may be overcome by use of type II MET TKIs (e.g. cabozantinib) (7–9), the genomic alterations favoring RAS-MAPK pathway activation described here will likely require a combination therapy strategy. In our *METex*14-mutated preclinical model system, RAS-MAPK pathway hyperactivation via *KRAS* overexpression or *NF1* downregulation induced MET TKI resistance that was overcome by the addition of the MEK inhibitor trametinib. Changes in the cfDNA profile suggested early evidence of molecular response to treatment with a crizotinib and trametinib combination therapy in a patient with acquired *KRAS* amplification, but treatment was poorly tolerated. Future efforts to develop combination therapies against these targets will require attention to agent selection, dosing, and scheduling to achieve both tolerability and efficacy.

This study enhances the understanding of the role of co-occurring genomic alterations in *METex*14-mutated NSCLC, with implications for the development of personalized therapeutic strategies to enhance the initial response magnitude and duration to MET TKI and delay or overcome acquired resistance to improve clinical outcomes. Given the prominence of genomic alterations favoring RAS-MAPK pathway hyperactivation, the addition of a MEK or potentially an ERK inhibitor to MET TKI therapy is a promising combination therapy strategy which warrants further prospective study.

## Methods

### Patients

Patient samples were obtained and analyzed in accordance with IRB-approved protocols. The cfDNA analysis included 332 consecutive samples from 289 patients with advanced (stage IIIB/IV) non-small cell lung cancer with a *METex*14 mutation obtained between October 2015 and March 2018, and 1653 consecutively tested samples from 1489 patients with *EGFR*-mutated NSCLC (L858R and del19) obtained between April 2016 and May 2017, as well as a previously published (11) cohort of TKI-naïve EGFR-mutated NSCLC.

### Cell lines and reagents

Ba/F3 cells were purchased from ATCC (ATCC^®^ HB-283^™^) and maintained in culture for a total of approximately 2-3 months in DMEM supplemented with 1 ng/mL IL-3 (Peprotech). Mycoplasma testing was not performed. Retrovirus was generated using TransIT-LT1 transfection reagent (Mirus). Cells were infected with filtered retrovirus, expressing either mCherry, human *MET^WT^* or human *MET^ex14^* in a pBABE-puro vector backbone as previously described (9) and selected in puromycin (2 μg/mL). Expression was confirmed by immunoblotting. KRAS-overexpressing cells were obtained by retroviral infection with a pBABE-hygro *KRAS4B* construct and selected in hygromycin (800 μg/mL). Knockdown of NF1 was achieved by lentiviral transduction of the following sequence: 5’-CCGGGCCAACCTTAACCTCTCTAATCTCGAGATTAGAGAGGTTAAGGTTGGCTTTTTG-3’, expressed from a pLKO.1-hygro plasmid backbone (Addgene #24150). Three days after lentiviral transduction, cells were selected via treatment with hygromycin B and knockdown was confirmed by immunoblotting. All drugs were purchased from Selleck Chemicals.

### Transformation and cell proliferation assays

Transformation assay was performed by removing IL-3 through centrifugation and adding 50 ng/mL human HGF (Peprotech 100-39H). For proliferation assays cells were seeded in 96-well plates at 5,000 cells/well and the following day were exposed to crizotinib (Selleck Chemicals, #S1068) at 0 to 10 μM and/or trametinib (Selleck Chemicals, #S2673) at 0.01 μM. After 72 hours of drug exposure, CellTiter-Glo (Promega) reagent was added and luminescence was measured on a Spectramax spectrophotometer (Molecular Devices, Sunnyvale, CA, USA) according the manufacturer’s instructions. Data are presented as percentage of viable cells compared with control cells (vehicle treatment).

### Immunoblotting

Cells were washed in PBS and lysed with 25 mM Tris-HCL (pH 7.6), 150 mM NaCl, 1% NP-40, 1% sodium deoxycholate, 0.1% SDS supplemented with Halt Protease Inhibitor Cocktail (Thermo Fisher Scientific) and Halt Phosphatase Inhibitor Cocktail (Thermo Fisher Scientific). Lysates were separated in a 4%–15% SDS-PAGE gel and transferred onto a nitrocellulose membrane (Bio-Rad). Membranes were blocked with 5% fetal bovine serum (FBS) in Tris-buffered saline (TBS) containing 0.1% Tween and incubated with the appropriate antibodies. Detection was performed via ECL Prime (Amersham Biosciences). Antibodies against the following were obtained from Cell Signaling Technology (Danvers, MA, USA) and were used at a dilution of 1:1000: MET (#3148), p-MET Y1349 (#3121), pMEK S217/221 (#9121), ERK1/2 (#3493), p-ERK1/2 T202/Y204 (#9106), HSP90 (#4874), PARP (#9546), NF1 (#14623), horseradish peroxidase (HRP)-conjugated anti-mouse (#7076) and HRP-conjugated anti-rabbit (#7074). The following antibody was obtained from EMD Millipore (Burlington, MA, USA): RAS (05-516, 1:2000 dilution). Detection was performed via ECL Prime (Amersham Biosciences).

### Cell-free DNA analysis

Samples were shipped to a Clinical Laboratory Improvement Act (CLIA)-certified, College of American Pathologists–accredited laboratory (Guardant Health, Redwood City, CA). cfDNA was extracted from whole blood collected in 10-mL Streck tubes. After double centrifugation, 5–30 ng of cfDNA was isolated for digital sequencing of either a 70 or 73-gene panel (Supplementary Table 6) as previously described (15). Only those genes in common to these two panels were included in subsequent analysis. Nonsynonymous mutations were further processed with the R statistical computing program (version 3.3). Variants with unknown or neutral predicted functional significance were filtered prior to analysis as previously described (11). Mutations previously reported as associated with clonal hematopoiesis were also excluded (16). Assignment as clonal or subclonal was performed by normalized mutational allele frequency to percentage detected using a cut off of 0.2 as previously described(11). Residue numbering was standardized to MET UniProtKB-P08581.

### Next Generation Sequencing

Tumor sample next generation sequencing (NGS) was performed in CLIA-approved laboratories. The Foundation One and Foundation ACT assays are commercially available assays which were used in the clinical standard-of-care setting. The UCSF500 assay sequences 479 cancer-associated genes to target 200X coverage, utilizing sequencing of a PBMC sample, target 100X coverage, as a control. The University of Florida GatorSeq NGS assay, utilized for both tumor tissue and PBMC analysis, performs sequencing of 76 cancer-associated genes with target 500X coverage (Supplemental Table 2). Germline mutations were subtracted utilizing sequencing of buccal swab samples to a target depth of 100X.

### Droplet Digital PCR

Isolated genomic DNA extracted from FFPE was amplified using ddPCR Supermix for Probes (Bio-Rad) using KRAS and MET assays (PrimePCR ddPCR Mutation Assay, Bio-Rad, and custom-designed). DNA template (8 μL) was added to 10 μL of ddPCR Supermix for Probes (Bio-Rad) and 2 μL of the primer/ probe mixture. This 20-μL reaction mix was added to a DG8 cartridge together with 70 μL of Droplet Generation Oil for Probes (Bio-Rad) and used for droplet generation. Droplets were then transferred to a 96-well plate (Eppendorf) and then thermal cycled with the following conditions: 5 minutes at 95°C, 40 cycles of 94°C for 30 seconds, 55°C for 1 minute followed by 98°C for 10 minutes (ramp rate 2°C/second). Droplets were analyzed with the QX200 Droplet Reader (Bio-Rad) for fluorescent measurement of FAM and HEX probes. Gating was performed based on positive and negative controls, and mutant populations were identified. The ddPCR data were analyzed with QuantaSoft analysis software (Bio-Rad) to obtain fractional abundance of the mutant DNA alleles in the wild-type (WT)/normal background. The quantification of the target molecule was presented as number of total copies (mutant plus WT) per sample in each reaction. Fractional abundance is calculated as follows: F.A. % = (Nmut/(Nmut + Nwt)) × 100), where Nmut is number of mutant events and Nwt is number of WT events per reaction. Multiple replicates were performed for each sample. ddPCR analysis of normal control genomic DNA (gene fragment obtained from IDT) and no DNA template (water) controls was performed in parallel with all samples, including multiple replicates as contamination-free controls.

### Statistical Analysis

Pairwise sample group comparisons for cfDNA analysis were carried out using a two-tailed Fisher’s exact t-test, with Benjamini-Hochberg correction for multiple comparisons, using a false discovery rate of less than 0.2. For cell viability curves, comparisons were performed using the two-sided student t-test, with significance threshold of *p*-value < 0.05.

## Acknowledgements

The authors acknowledge funding support from NIH / NCI U54CA224068 (to R.B.C.); R01CA227807, R01CA239604, R01CA230263 (to E.A.C.); NIH / NCI U01CA217882, NIH / NCI U54CA224081, NIH / NCI R01CA204302, NIH / NCI R01CA211052, NIH / NCI R01CA169338, and the Pew-Stewart Foundations (to T.G.B.); and the Damon Runyon Cancer Research Foundation, Doris Duke Charitable Foundation, V Foundation, and American Cancer Society to C.M.B.

