## Supplemental Figures and Tables for "Co-occurring alterations in the RAS-MAPK pathway limit response to MET inhibitor treatment in *MET* exon 14 skipping mutation positive lung cancer"

**Supplemental Table 1**

| | ***MET*ex14 Cohort** | ***EGFR*m Cohort** |
|---|---|---|
| Total # Samples | 332 | 1653 |
| Total # Patients | 289 | 1489 |
| Date Range | 10/2015-3/2018 | 4/2016-5/2017 |
| Gender (% Patients) | | |
| Female | 172 (59.5%) | 992 (66.6%) |
| Male | 116 (40.1%) | 497 (33.4%) |
| Not Specified | 1 (0.3%) | 0 (0%) |
| Mean Age | 73 years | 64.4 years |
| Stage III/IV (% Samples) | 289 (100%) | 1653 (100%) |
| Histology (% Patients) | | |
| Lung Adenocarcinoma | 163 (56.4%) | 532 (35.7%) |
| NSCLC, not otherwise specified | 93 (32.2%) | 954 (64.1%) |
| Lung Squamous Cell Carcinoma | 20 (6.9%) | 0 (0%) |
| Other | 13 (4.5%)[a] | 3 (0.2%)[b] |
| Treatment History (% Samples) | | |
| Unknown | 261 (78.6%) | 1653 (100%) |
| Tyrosine Kinase Inhibitor[c] | 26 (7.8%) | |
| Chemotherapy[d] | 22 (6.6%) | |
| Checkpoint Inhibitor | 17 (5.1%) | |
| Radiation | 3 (0.9%) | |

[a]Lung cancer NOS (2%), Carcinoid (0.7%), Sarcomatoid (0.7%), Large Cell (0.3%), Carcinosarcoma (0.3%), Small Cell Lung Cancer (0.3%)
[b]Lung cancer, not otherwise specified
[c]Crizotinib (3.3%), Erlotinib (2.7%), Alectinib (0.6%), MET TKI not otherwise specified (0.3%), Crizotinib/trametinib (0.3%), Cabozantinib (0.3%), Afatinib (0.3%)
[d]Including chemotherapy combinations: Pemetrexed/Pembrolizumab (0.3%), docetaxel/nintedanib (0.3%)

**Supplemental Table 1. Cell-free DNA NSCLC clinical cohort demographic and clinical information.** Clinical and demographic data for two cohorts of patients with advanced NSCLC, incorporating all patients with either the *MET* exon 14 skipping mutation (*MET*ex14) or an activating epidermal growth factor mutation (*EGFR*m) identified by cfDNA sequencing over the specified time period.

### Supplemental Table 2

| NGS Panel | Gene List |
|---|---|
| **Guardant360, 70-Gene Assay** | AKT1, ALK, APC, AR, ARAF, ARID1A, ATM, BRAF, BRCA1, BRCA2, CCND1, CCND2, CCNE1, CDH1, CDK4, CDK6, CDKN2A, CTNNB1, EGFR, ERBB2, ESR1, EZH2, FBXW7, FGFR1, FGFR2, FGFR3, GATA3, GNA11, GNAQ, GNAS, HNF1A, HRAS, IDH1, IDH2, JAK2, JAK3, KIT, KRAS, MAP2K1, MAP2K2, MET, MLH1, MPL, MYC, NF1, NFE2L2, NOTCH1, NPM1, NRAS, NTRK1, PDGFRA, PIK3CA, PTEN, PTPN11, RAF1, RB1, RET, RHEB, RHOA, RIT1, ROS1, SMAD4, SMO, STK11, TERT, TP53, TSC1, VHL, CDKN2B[a], SRC[a] |
| **Guardant360, 73-Gene Assay** | AKT1, ALK, APC, AR, ARAF, ARID1A, ATM, BRAF, BRCA1, BRCA2, CCND1, CCND2, CCNE1, CDH1, CDK4, CDK6, CDKN2A, CTNNB1, DDR2[a], EGFR, ERBB2, ESR1, EZH2, FBXW7, FGFR1, FGFR2, FGFR3, GATA3, GNA11, GNAQ, GNAS, HNF1A, HRAS, IDH1, IDH2, JAK2, JAK3, KIT, KRAS, MAP2K1, MAP2K2, MAPK1[a], MAPK3[a], MET, MLH1, MPL, MTOR[a], MYC, NF1, NFE2L2, NOTCH1, NPM1, NRAS, NTRK1, NTRK3[a], PDGFRA, PIK3CA, PTEN, PTPN11, RAF1, RB1, RET, RHEB, RHOA, RIT1, ROS1, SMAD4, SMO, STK11, TERT, TP53, TSC1, VHL |
| **University of Florida GatorSeq NGS Assay** | ABL1, AKT1, ALK, ASXL1, BAALC, BCOR, BCR, BRAF, BRINP3, CBFB, CEBPA, CRLF2, CTNNB1, DDR2, DEK, DNMT3A, EGFR, ERBB2, ERG, ETV6, EZH2, FBXW7, FGFR1, FGFR2, FLT3, GNA11, GNAQ, HOXA9, HRAS, IDH1, IDH2, JAK2, KIT, KMT2A, KRAS, MAP2K1, MECOM, MET, MKL1, MLLT3, MN1, MPL, MYC, MYH11, NF1, NOTCH1, NPM1, NRAS, NUP214, PDGFRA, PHF6, PIK3CA, PML, PTEN, PTPN11, RAD21, RARA, RBM15, RET, RPN1, RUNX1, RUNX1T1, SF3B1, SMAD4, SMC1A, SMC3, SMO, SRSF2, STAG2, TET2, TP53 TSC1, U2AF1, U2AF2, WT1, ZRSR2 |
| **UCSF500 NGS Assay** | ABL1, ABL2, ACVR1, ACVR1B, AJUBA, AKT1, AKT2, AKT3, ALK, AMER1, APC, APOBEC3G, AR, ARAF, ARFRP1, ARHGAP35, ARID1A, ARID1B, ARID2, ARID5B, ASH2L, ASXL1, ASXL2, ATF1, ATM, ATR, ATRX, AURKA, AURKB, AXIN1, AXIN2, AXL, BAP1, BARD1, BCL2, BCL2A1, BCL2L1, BCL2L12, BCL2L2, BCL6, BCOR, BCORL1, BLM, BRAF, BRCA1, BRCA2, BRD4, BRIP1, BTG1, BTK, C11orf30, CALR, CARD11, CBFB, CBL, CBLB, CCND1, CCND2, CCND3, CCNE1, CD274, CD79A, CD79B, CDC42, CDC73, CDH1, CDK12, CDK4, CDK6, CDK8, CDKN1A, CDKN1B, CDKN2A, CDKN2B, CDKN2C, CEBPA, CHD1, CHD2, CHD4, CHD5, CHEK1, CHEK2, CIC, CLDN18, CNOT3, COL1A1, COL2A1, CRCT1, CREB1, CREBBP, CRKL, CSF1R, CSF3R, CTCF, CTNNA1, CTNNB1, CUL3, CUX1, CXCR4, CYLD, DCC, DDIT3, DDR2, DDX3X, DDX41, DGKH, DICER1, DIS3, DNAJB1, DNMT3A, DOT1L, DUSP2, DUSP4, DUSP6, DYNC1I1, EBF1, EDNRB, EGFR, EGR1, EIF1AX, ELF3, EP300, EPCAM, EPHA2, EPHA3, EPHA5, EPHA7, EPHB1, EPOR, ERBB2, ERBB3, ERBB4, ERCC1, ERCC2, ERG, ERRFI1, ESPL1, ESR1, ESR2, ETS1, ETV6, EWSR1, EZH1, EZH2, FAM46C, FANCA, FANCC, FANCE, FANCF, FANCG, FANCL, FAT1, FAT3, FBXW7, FGF10, FGF14, FGF19, FGF23, FGF3, FGF4, FGF6, FGFR1, FGFR2, FGFR3, FGFR4, FH, FLCN, FLT1, FLT3, FLT4, FOXA1, FOXL2, FOXO1, FOXP1, FRS2, FUBP1, FUS, FYN, GAB2, GATA1, GATA2, GATA3, GLI1, GLI2, GNA11, GNA13, GNAQ, GNAS, GPC3, GPR124, GRIN2A, GRM3, GSK3B, H3F3A, H3F3B, HDAC4, HDAC9, HEY1, HGF, HIF1A, HIST1H3B, HMGA2, HNF1A, HOXB13, HRAS, HSP90AB1, HSPA2, HSPA5, ID3, IDH1, IDH2, IGF1R, IGF2, IGF2R, IKBKE, IKZF1, IKZF2, IKZF3, IL2RB, IL7R, INHBA, INPP4B, IPMK, IRF4, IRS2, JAK1, JAK2, JAK3, JAZF1, KAT6A, KDM5A, KDM5C, KDM6A, KDR, KEAP1, KIT, KLF4, KLHL6, KMT2A, KMT2B, KMT2D, KNSTRN, KRAS, LEF1, LIFR, LRP1B, LZTR1, MALAT1, MAML2, MAP2K1, MAP2K2, MAP2K4, MAP3K1, MAP3K2, MAP3K5, MAP3K7, MAP3K9, MAPK1, MCL1, MDM2, MDM4, MED12, MEF2B, MEN1, MET, MGA, MGMT, MITF, MLH1, MLH3, MPL, MRE11A, MSH2, MSH3, MSH6, MTOR, MUTYH, MYB, MYBL1, MYC, MYCL, MYCN, MYD88, MYH9, NAV3, NBN, NCKAP5, NCOA2, NCOA3, NCOR1, NF1, NF2, NFE2L2, NFKBIA, NFKBIE, |

| | |
|---|---|
| | NIPBL, NKX2-1, NOTCH1, NOTCH3, NPM1, NRAS, NSD1, NT5C2, NTRK1, NTRK2, NTRK3, NUP93, NUTM1, OR5L1, PAK1, PAK3, PALB2, PARK2, PAX3, PAX5, PAX7, PAX8, PBRM1, PDCD1LG2, PDGFB, PDGFRA, PDGFRB, PDK1, PHF6, PHOX2B, PIK3CA, PIK3CG, PIK3R1, PIK3R2, PLAG1, PLCB4, PMS1, POLD1, POLE, POLQ, POT1, POU3F2, PPM1D, PPP2R1A, PPP6C, PRDM1, PREX2, PRKACA, PRKAG2, PRKAR1A, PRKCA, PRKCH, PRKDC, PTCH1, PTCH2, PTEN, PTK2B, PTPN1, PTPN11, PTPRB, PTPRD, PTPRK, PTPRT, RAC1, RAD21, RAD50, RAD51, RAD51C, RAD51D, RAF1, RARA, RASA1, RASA2, RB1, RBM10, REL, RELA, RET, RHEB, RHOA, RICTOR, RIT1, RNF43, ROBO1, ROS1, RPL10, RPTOR, RRAGC, RRAS, RRAS2, RSPO2, RSPO3, RUNX1, RUNX1T1, SDHB, SDHD, SETBP1, SETD2, SF3B1, SH2B3, SHH, SIN3A, SLIT2, SLITRK6, SMAD2, SMAD3, SMAD4, SMARCA2, SMARCA4, SMARCB1, SMC1A, SMC3, SMO, SNCAIP, SOCS1, SOS1, SOS2, SOX10, SOX2, SOX9, SPEN, SPOP, SPRED1, SPRY1, SPRY2, SPRY4, SPTA1, SRC, SRSF2, SS18, STAG2, STAT3, STAT4, STAT6, STK11, SUFU, SYK, SYNE1, TADA1, TBX3, TCEB1, TCF7L2, TERT, TET2, TFE3, TFEB, TGFBR2, TLR4, TMPRSS2, TNFAIP3, TNFRSF14, TOP1, TOP2A, TP53, TRAF3, TRAF7, TRIM28, TSC1, TSC2, TSHR, TSHZ2, TSHZ3, TSLP, TTYH1, TYK2, U2AF1, USP7, VEGFA, VHL, WHSC1, WISP3, WRN, WT1, XBP1, XPO1, YAP1, YWHAE, ZBTB20, ZFHX3, ZFHX4, ZMYM3, ZNF217, ZNF703, ZRSR2 |

[a]Excluded from 68 gene set in common between 70 and 73 gene sets

**Supplemental Table 2. Genes included in NGS Panels.** List of cancer-associated genes included in the 70- and 73-gene versions of the Guardant360 assay, University of Florida GatorSeq NGS assay, and UCSF500 NGS assay.

### Supplemental Table 3

| Patient # | Histology | Sample Prior to Known MET TKI Exposure | | Sample After MET TKI Exposure | |
|---|---|---|---|---|---|
| | | Treatment Status | Genomic Alterations | MET TKI Received | Genomic Alterations % cfDNA/CNG |
| 1 | Lung Adeno-carcinoma | Pre-treatment | METex14 (c.3028+3A>G, 7.6%), PTEN E43Q (0.1%), CCNE1 CNG (2.3) | Crizotinib | METex14 (0.9-5.1%), BRAF R199G (0.1-0.2%), CCNE1 CNG (2.2), PIK3CA R38H (0.1%), MET Y1230S (0.3-1.5%), MET F1200I (0.2%), KRAS CNG (2.25-2.28) |
| 2[a] | Lung Adeno-carcinoma | Pre-treatment | METex14 (c.2888-19_2895del27, 8%)[b] | Crizotinib | METex14 (2.2%), KRAS G12D (7.8%) |
| 3 | Lung Adeno-carcinoma | Carboplatin/ pemetrexed/ bevacizumab | METex14 (c.3028G>C, 1.2%), BRCA1 R691G (0.1%) | Crizotinib | METex14 (1%), FBXW7 R689Q (0.1%), AR W742C (0.1%), TP53 Y163S (0.1%), KIT CNG (2.2) |
| 4 | Lung Adeno-carcinoma | Pre-treatment | METex14 (c.3028G>C, 19.8%), NF1 I1499V (0.3%), BRAF CNG (2.6), NF1 R1534Q (0.1%), EGFR V851A (0.1%), EGFR CNG (2.5), MET CNG (2.6), CDK6 CNG (2.6), MYC CNG (2.4) | Crizotinib[c] | METex14 (2.5%), NF1 I1499V (0.2%), CCND1 CNG (2.5), TP53 N239S (1.8%), TP53 R248W (0.5%), NF1 p.Gln2636fs (1.1%), MET p.Ser244fs (1.2%) |
| 5 | Lung Adeno-carcinoma | Pre-treatment | METex14 (c.3028+1delG), MET CNG, CDK6 CNG, ATM splice site, MDM2 CNG, CDKN2A/B loss[b] | Crizotinib | METex14 (53.4%), MET CNG (3.9), CDK6 CNG (3), AR CNG (2.3), PIK3CA CNG (2.3), MET Y1230H (9.1%), MET D1228N (4.3%), EGFR CNG (2.2) |
| | | | | Glesatinib | METex14 (63.5%), MET CNG (4.9), CDK6 CNG (3.3), PIK3CA CNG (2.4), MET L1195V (0.8%), MET D1228N (13.9%) |
| 6 | Lung SCC | Nivolumab | METex14 (c.2888-12_2889delCTCTGTTTTA AGATinsTAAGAG, 4.6%) | Crizotinib | METex14 (7.7%), EGFR CNG (2.7), TP53 p.Arg156del (0.04%) |
| 7 | Lung Adeno-carcinoma | Carboplatin/ Paclitaxel | TP53 P27L (0.2%), TP53 c375+1G>C (0.1%) | Crizotinib | METex14 (c.2888-20_2888delTTCTTTCTCT CTGTTTTAAGA, 0.02%) |
| 8 | Lung Adeno-carcinoma | Carboplatin/ pemetrexed/ bevacizumab | METex14 (c.3028+2T>C, 0.4%), BRAF S273G (0.5%), MET R1170* (47.7%) | Crizotinib | BRAF S273G (0.3%), TP53 V173M (0.3%) |
| 9 | Lung Adeno-carcinoma | Unknown | METex14 (c.3028G>T, 14.5%), MET CNG (2.4), EGFR K80T (0.8%), ERBB2 N68S (0.7%), KRAS G12S (0.1%) | Crizotinib | METex14 (50%), MET CNG (4), EGFR K80T (0.3%), ERBB2 N68S (3.5%), MET L1195V (16.6%), TP53 V216E (0.1%), EGFR CNG (2.4) |

| | | | | | |
|---|---|---|---|---|---|
| 10 | Lung Adeno-carcinoma | Unknown | METex14 (c.3012_3028+3delAGCTACTTTTCCAGAAGGTAinsG, 4.8%), MET CNG (2.2), CDKN2A p.Thr77fs (0.9%) | Crizotinib | TP53 R158H (0.2%), EGFR R836H (0.2%), PDGFRA R558H (0.2%) |
| 11 | NSCLC NOS | Pre-treatment | METex14 (c.2888-5_2905delTTAAGATCTGGGCAGTGAATTA, 2%), RECQL4 splice site (2464-1G>C, 46%), SMAD4 Q224X (10%), CDK4 CNG (22.4), KMT2A CNG (7.2), MDM2 CNG (11.5)[b] | Crizotinib | METex14 (0.6%), MET D1228H (0.1%), TP53 F270L (0.1%) |
| 12 | Lung Adeno-carcinoma | Pre-treatment | METex14 (c.2888-20_2898del13), ERBB4 E69K, CDK4 CNG, GLI1 CNG, MDM2 CNG, APC E1284K[b] | Crizotinib | METex14 (3.3%), ATM N3003T (11%) |

[a]cfDNA analysis prior to crizotinib via Foundation ACT assay also notable for KRAS G12D which was not detected upon sequencing (University of Florida in-house NGS assay) of a pre-treatment tumor biopsy sample. Sequencing of a tumor biopsy of a differing progressing site following crizotinib treatment (Foundation One) was notable for both *KRAS* G12D and *KRAS* amplification.
[b]Sequencing prior to known MET TKI exposure performed on a tumor biopsy sample rather than via plasma cfDNA analysis, Foundation One (patients 5 and 13), or Cancer-Select assay (patient 12).
[c]Followed by pemetrexed prior to cfDNA testing.

**Supplemental Table 3. Genomic alterations in the cfDNA of patients treated with a MET TKI.** The genomic alterations newly identified upon targeted sequencing for cancer-associated genes in cfDNA samples obtained following known MET TKI exposure compared to results of samples obtained prior to known MET TKI exposure. Sequencing performed via the Guardant 360 assay, unless otherwise specified. Further details for patient one and two in Supplemental Figure 2. Additional details regarding patient five have previously been published (7,9). Abbreviations: CNG, copy number gain; NOS, not otherwise specified; SCC, squamous cell carcinoma.

### Supplemental Table 4

| Kinase | Residue | Amino Acid Sequence | Reported Mutation | |
|---|---|---|---|---|
| **MET** | F1200 | KYLASKK**F**VHRDLAARNCML | F1200I | Type II > Type I MET TKI resistance predicted by *in vitro* studies(21) |
| **ALK** | F1245 | QYLEENH**F**IHRDIAARNCLL | F1245C | Intermediate crizotinib resistance(22)<br>ALK-activating mutation(23) |
| **ROS1** | F2075 | VYLERMH**F**IHRDLAARNCLV | F2075V | Cabozantinib resistance(24) |
| **ABL** | F359 | EYLEKKN**F**IHRDLAARNCLV | F359V | Imatinib and nilotinib resistance(25,26) |
| **NTRK1** | F646 | VYLAGLH**F**VHRDLATRNCLV | F646I | Cabozantinib resistance(27) |

**Supplemental Table 4. The *MET* F1200 residue is conserved across multiple tyrosine kinases.** Sequence alignment was performed utilizing protein BLAST (NCIB)(43), demonstrating conserved residues (shown in yellow) corresponding to *MET* F1200 in *ALK*, *ROS1*, *ABL*, and *NTRK1* with associated influence on TKI response to the type II TKIs cabozantinib, imatinib, and nilotinib, and to the type I TKI crizotinib.

**Supplemental Figure S1**

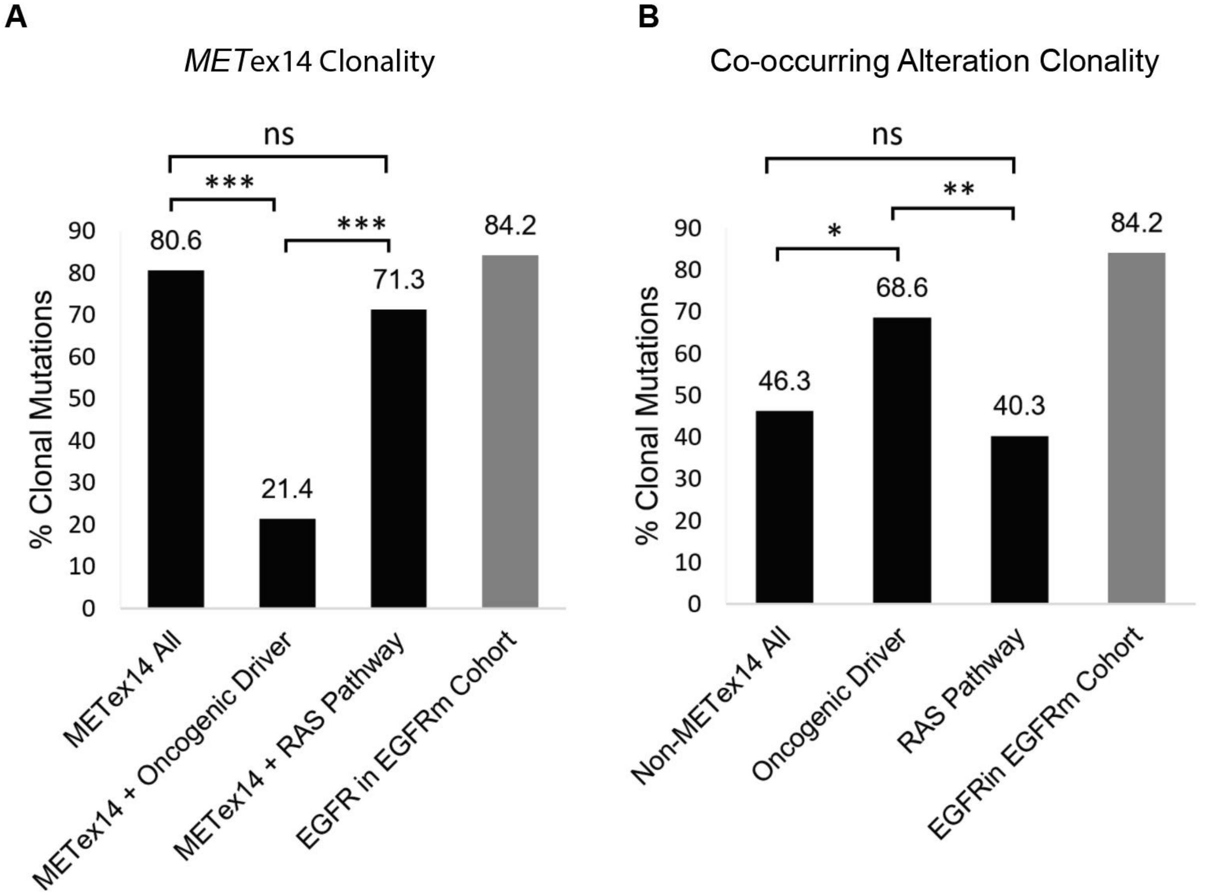


**Supplemental Figure S1. Clonality of *MET*ex14 mutations and of detectable co-occurring mutations within the cfDNA of patients with *MET*ex14-mutated NSCLC.** A. Clonality of the *MET*ex14 skipping mutation in the whole cohort (n=289 mutations), and in samples with a co-occuring canonical oncogenic driver (n=28) or a co-occurring RAS pathway alteration (n=94). B. Clonality of detectable co-occurring genomical alterations in the cfDNA of the same cohort of pateints with *MET*ex14-mutated NSCLC for all co-occuring alterations (n=825), co-occurring canonical onocgenic driver mutations (n=35), or co-occurring RAS pathway alteartions (n=129). In both panels the clonality of canonical EGFR mutations (n=1645, *EGFR* del19, *EGFR* L858R, and *EGFR* T790M) from an independent cohort of patients with *EGFR*-mutated (*EGFR*m) NSCLC is displayed (light gray) for comparison. *** p-value < 0.001, ** p-value < 0.01, * p-value < 0.05

**Supplemental Figure S2**

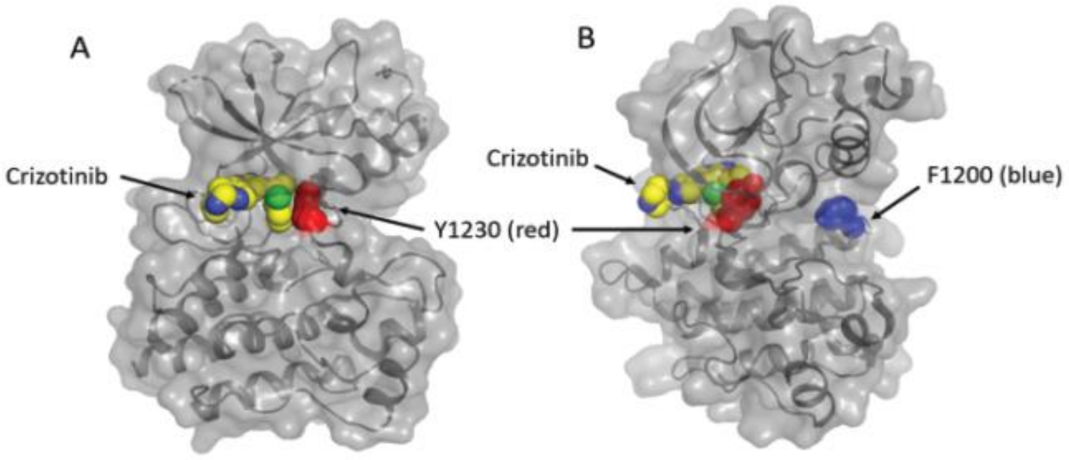


**Supplemental Figure S2. MET tyrosine kinase domain modeling location of the F1200 residue.** *MET* Y1230 and F1200 residues modeled in their structural context based on PDB 2WGJ. **A.** Crizotinib is shown in yellow, in close approximation to the *MET* Y1230 residue (red), as a frequently reported site of type I MET TKI resistance mutations. **B.** The location of the F1200 mutation at the DFG-out pocket, at a disparate location from the crizotinib binding site, is highlighted in blue.

### Supplemental Figure S3

**A**

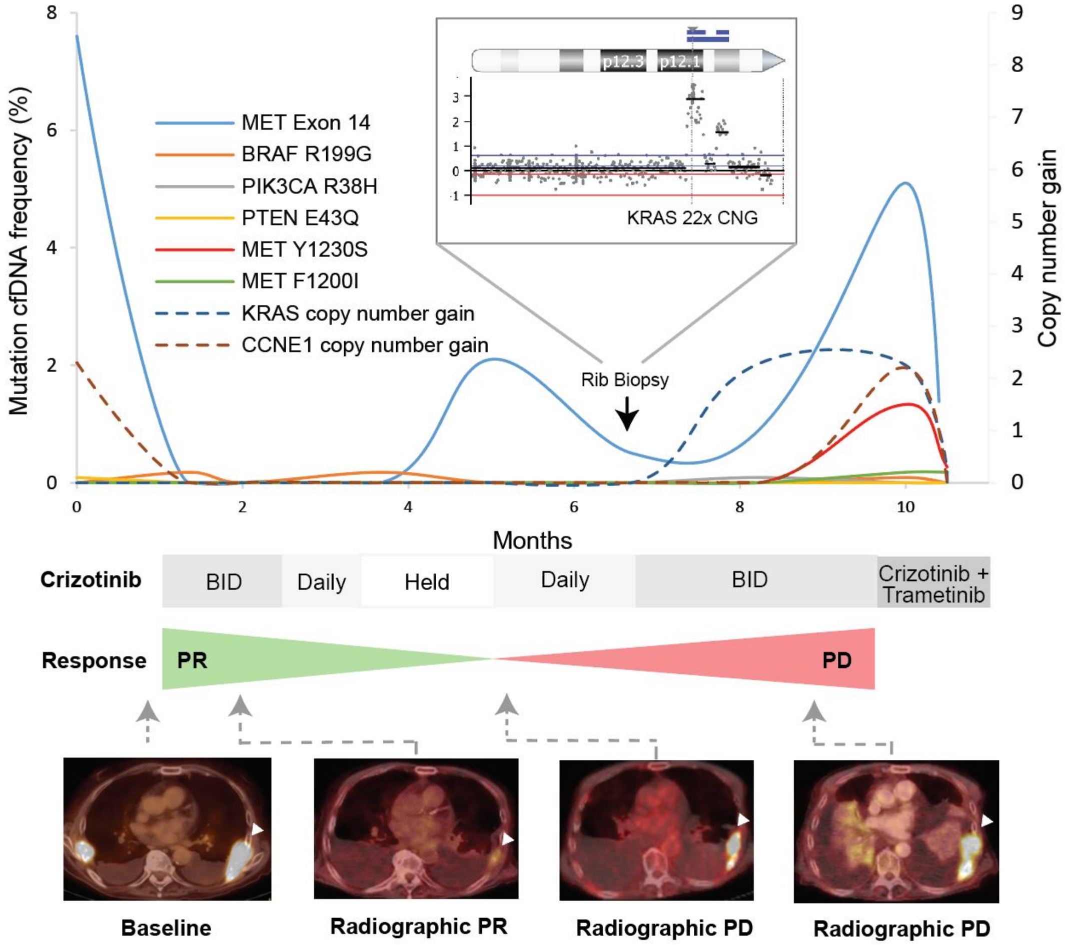


**B**

| | RUL Lung VAF(%)/CNG† | Plasma cfDNA Allele Frequency (%)‡ | Abdominal Lesion VAF(%)/CNG* |
|---|---|---|---|
| **MET Exon14** | 8% | 2.2% | 77.9% |
| **KRAS** | Wild type | G12D 7.8% | G12D 47.4%, 11x CNG |
| **ddPCR** | METex14 11%<br>KRAS G12D not detected | | |

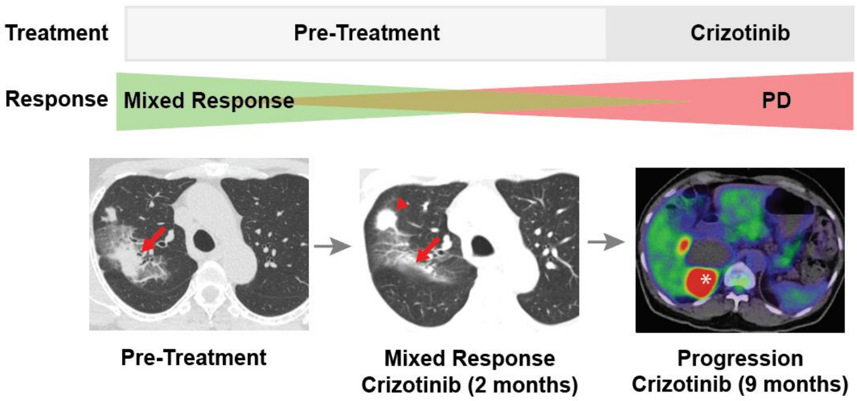

**Supplemental Figure S3. *KRAS* amplification and/or *KRAS G12D* mutation in *MET*ex14-mutated NSCLC with resistance to crizotinib. A.** Serial cfDNA analysis for a targeted panel of cancer-associated genes (Supplemental Table S6) in a patient with stage IV MET exon 14 mutated NSCLC, obtained prior to treatment, during partial response to crizotinib and through the development of acquired resistance to crizotinib treatment. The timing of a biopsy of a crizotinib-resistant left-sided metastatic rib lesion, on which a UCSF clinical NGS panel was performed, is shown (black arrow) as are representative PET/CT images, including the sampled left-sided rib metastasis (white arrow head). The inset displays copy number variation at the short arm of chromosome 12 as determined by CNVkit (42) analysis of UCSF500 assay data obtained from a metastatic soft tissue rib lesion, showing 22-fold KRAS amplification. **B.** Serial targeted DNA sequencing of cancer-associated genes performed on tumor tissue samples or plasma cfDNA in a patient with advanced-stage *MET*ex14-mutated NSCLC. DNA sequencing was performed on a pre-treatment biopsy of a lung lesion (red arrow) using a clinical University of Florida gene panel assay and via a commercial plasma cfDNA panel (Foundation ACT) prior to mixed response to crizotinib treatment (response at original lesion, progression at prior small right upper lobe lung lesion highlighted by a red arrowhead). This sequencing demonstrated a *METex14* mutation (2888-19_2895del27), with wild-type *KRAS* in the tumor biopsy sample, while cfDNA testing demonstrated the known *MET*ex14 mutation and an activating *KRAS* G12D mutation. Confirmatory droplet digital PCR (ddPCR) testing verified absence of the *KRAS* G12D mutation within the pre-treatment biopsy sample to a sensitivity of <0.02% Tissue NGS testing on a tissue sample of a progressing abdominal lesion (white asterisk) at acquired crizotinib resistance was notable for the known *MET*ex14 mutation (77.9% VAF) and *KRAS* G12D mutations (47.4% VAF), and also found *KRAS* copy number gain (~11-fold), along with other gene alterations of less certain significance (Supplemental Table S5). Next-generation DNA sequencing of the patient's peripheral blood mononuclear cells (PBMCs) confirmed absence of detectable *KRAS* G12D mutation within the hematopoietic lineage.

### Supplemental Figure S4

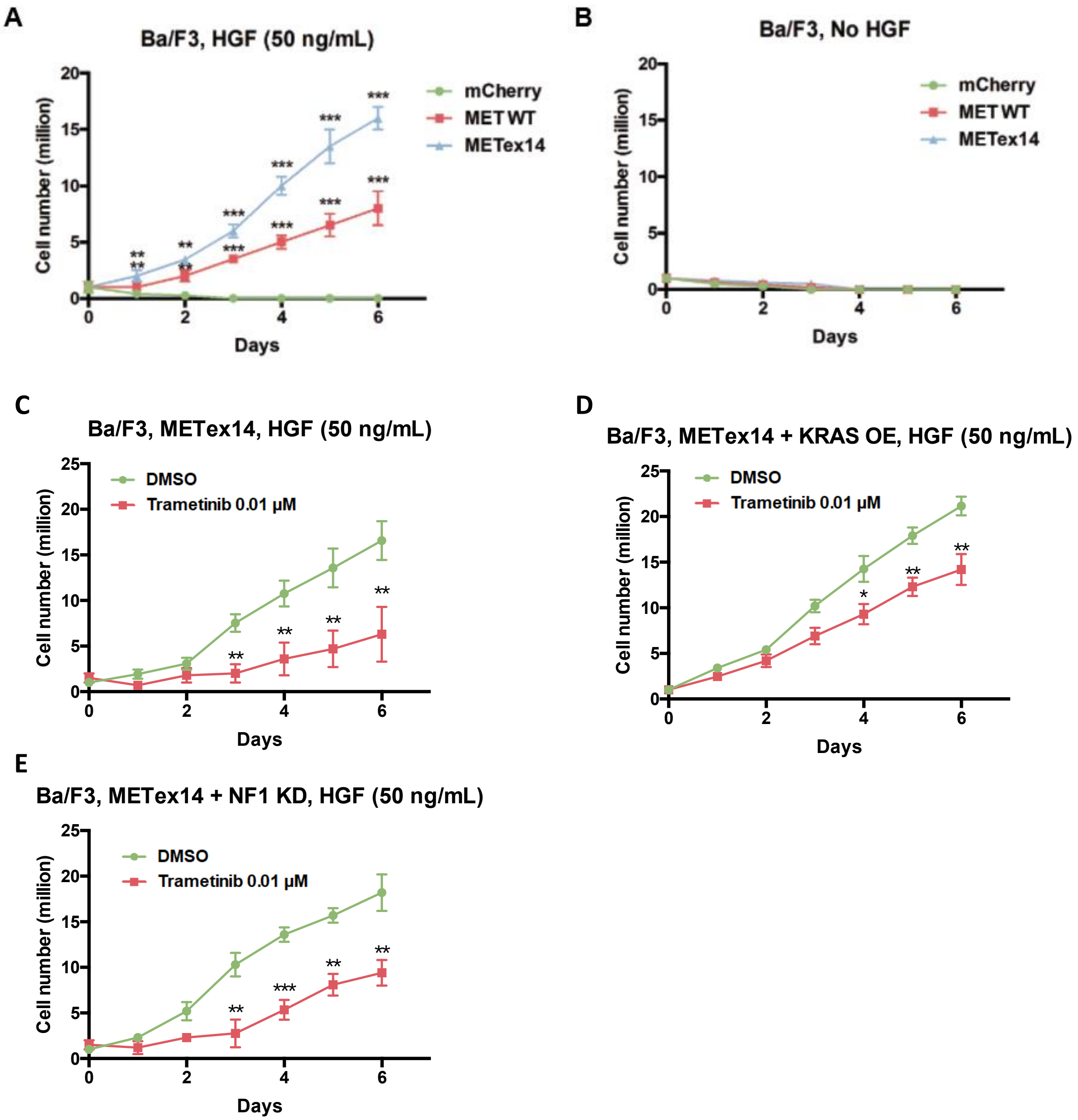


**Supplemental Figure S4. Relative cell viability of Ba/F3 *MET*ex14-mutant expressing cells treated with trametinib monotherapy** **A.** Ba/F3 cells with wild type *MET*, *MET*ex14, or mCherry control demonstrating acquisition of IL-3 independent growth with addition of HGF (50 ng/mL) in cells expressing the *MET* exon 14 mutant and to a lesser extent in cells expressing wild type *MET*. **B.** Cell growth curves of Ba/F3 cells expressing wild type *MET*, *MET*ex14, or mCherry control without growth in the absence of IL-3 and HGF. **C.** Ba/F3 cells overexpressing *MET*ex14 were treated with HGF (50 ng/mL) with either DMSO control or trametinib 0.01 µM.  Cell growth is reduced but not eliminated at the 0.01 µM dose of trametinib. **D.** Ba/F3 cells overexpressing both *MET*ex14 and *KRAS* were treated with HGF (50 ng/mL) with either DMSO control or trametinib 0.01 µM. **E.** Ba/F3 cells overexpressing *MET*ex14 and with shRNA knockdown of *NF1* were treated with HGF (50 ng/mL) with either DMSO control or trametinib 0.01 µM.  ** $p$-value < 0.01, *** $p$-value <0.001 by student's t-test.
